# Mechanistic Investigation of LncRNA-ZMIZ1-AS1 in the Malignant Phenotype of Hepatocellular Carcinoma

**DOI:** 10.1101/2025.11.23.690041

**Authors:** Guo Chen, Hao Tang, Chunhu Mao, Gang Wu

**Affiliations:** Department of General Surgery Deyang Hospital, Affiliated Hospital of Chengdu University of Traditional Chinese Medicine, Deyang 618000, China; Department of General Surgery, Sichuan Tianfu New Area People’s Hospital, Chengdu 610213, China; .Nanchong Hospital of Beijing Anzhen Hospital CMU, Nanchong 637000, China; Department of hepatopancreatobiliary Surgery, Sichuan Provincial People’s Hospital, University of Electronic Science and Technology of China,Chengdu 610072, China

**Keywords:** Hepatocellular carcinoma, LncRNA-ZMIZ1-AS1, PTBP1, NOTCH signaling pathway

## Abstract

**Background:** Hepatocellular carcinoma (HCC) is one of the most fatal cancers worldwide, with high recurrence and mortality rates. LncRNA-ZMIZ1-AS1 has attracted considerable attention for its role in the tumor microenvironment, particularly in the regulation of HCC cell growth, invasion, and biological behavior. However, the specific mechanisms of LncRNA-ZMIZ1-AS1 in HCC remain unclear.

**Methods:** We analyzed the relationship between LncRNA-ZMIZ1-AS1 and HCC clinical prognosis using bioinformatics with GEO and TCGA datasets. We also conducted molecular biology experiments, including cell culture, real-time PCR, and Western blotting, as well as cell viability and flow cytometry analyses, to investigate the effects of LncRNA-ZMIZ1-AS1 knockdown on HCC cell proliferation, migration, invasion, and apoptosis. Furthermore, we used Gene Set Enrichment Analysis (GSEA) to identify potential signaling pathways involving LncRNA-ZMIZ1-AS1.

**Results:** LncRNA-ZMIZ1-AS1 was highly expressed in HCC patients and significantly associated with clinical characteristics, such as gender and clinical stage. Knockdown of LncRNA-ZMIZ1-AS1 notably inhibited HCC cell proliferation, migration, and invasion while increasing apoptosis rates. Additionally, GSEA suggested a potential association between high LncRNA-ZMIZ1-AS1 expression and the NOTCH signaling pathway.

**Conclusion:** The high expression of LncRNA-ZMIZ1-AS1 in HCC likely promotes malignant phenotypes by regulating signaling pathways such as NOTCH. This finding underscores its potential as a biomarker for HCC, providing a theoretical basis for further research on its molecular mechanisms and clinical applications in liver cancer.

## Introduction

Hepatocellular carcinoma (HCC) is one of the most common fatal cancers, ranking among the highest in incidence and mortality. Hepatocellular carcinoma (HCC), often referred to simply as liver cancer, is the predominant type. Despite rapid advances in liver surgery, the long-term outcomes for HCC remain unsatisfactory, as tumor recurrence and metastasis are frequently observed. The five-year survival rate remains extremely low, with over 700,000 liver cancer-related deaths worldwide each year^[1]^. Clinically, only about 20% of new HCC cases qualify for curative resection; however, 70% of patients experience recurrence post-surgery, and 30% succumb to liver cancer within five years^[2–4]^. Accurate prognostic assessments are crucial in guiding treatment decisions for HCC patients, underscoring the urgent need for novel biomarkers to predict HCC prognosis and reveal therapeutic targets.

Non-coding RNAs (ncRNAs) are RNA molecules that do not encode proteins but play essential roles in gene regulation, cellular differentiation, and development. NcRNAs can be categorized into several types, including small ncRNAs (e.g., miRNA and siRNA), long non-coding RNAs (lncRNAs), and circular RNAs (circRNAs)^[5–7]^. NcRNAs influence gene expression through various mechanisms. For example, small ncRNAs typically regulate gene expression at the post-transcriptional level by binding to the 3’ untranslated region of mRNAs, thereby inhibiting expression. Conversely, lncRNAs can regulate transcription by providing structural support, acting as molecular “sponges” to absorb other small RNA molecules, or directly interacting with proteins to influence the transcriptional process^[8, 9]^.

LncRNA-ZMIZ1-AS1, a type of long non-coding RNA, functions primarily through interactions with other molecules, including proteins, small RNAs, or DNA. It may regulate gene expression by impacting transcriptional regulation, mRNA stability, or protein translation. In cancer biology, LncRNA-ZMIZ1-AS1 may contribute to cancer initiation, progression, and metastasis by modulating intercellular communication in the tumor microenvironment, influencing tumor cell proliferation and apoptosis, or engaging in tumor metabolism. Specifically, LncRNA-ZMIZ1-AS1 can regulate the stability of ZMIZ1 mRNA by interacting with the RNA-binding protein PTBP1. ZMIZ1-AS1 recruits PTBP1 to maintain ZMIZ1 expression, thereby promoting growth, invasion, and migration in osteosarcoma cells, a classic function of lncRNAs in modulating the expression of adjacent genes^[10]^.

Research teams have explored the expression and function of lncRNA ZMIZ1-AS1 in glioblastoma, where it was observed that ZMIZ1-AS1 expression was significantly elevated and closely associated with the malignant phenotype. Knockdown of ZMIZ1-AS1 led to reduced tumor cell proliferation, invasion, and migration, while apoptosis rates increased. Further molecular studies revealed that ZMIZ1-AS1 regulates the malignant phenotype of glioblastoma cells by modulating ZMIZ1 expression through its interaction with PTBP1. These findings indicate that ZMIZ1-AS1 plays a critical role in the pathological processes of both osteosarcoma and glioblastoma.

In studies related to HCC, it was found that circZMIZ1 promotes HCC progression by suppressing CD8+ T cell antitumor activity through modulation of the miR-15a-5p/KCNJ2 axis. This suggests a potential link between ZMIZ1-AS1 expression, tumor immune evasion, and immunotherapy effectiveness^[11]^. Additionally, JIANG^[12]^ et al. reported that circZMIZ1 expression was elevated in the plasma of prostate cancer patients compared to those with benign prostatic hyperplasia. Knockdown of circZMIZ1 in prostate cancer cells inhibited proliferation and caused G1 phase cell cycle arrest. Mechanistically, the authors suggested that circZMIZ1 promotes cancer development by increasing the expression of androgen receptor (AR) and its splice variant AR-V7. However, there are no existing studies on LncRNA-ZMIZ1-AS1 in HCC, highlighting the necessity of exploring and validating its potential role in this type of cancer.

## Materials and Methods

### Bioinformatics Analysis

Based on TCGA and GEO ^[13, 14]^data cohorts, we utilized Pearson correlation analysis in R to evaluate the correlation between m6A-modifying factors and LncRNA-ZMIZ1-AS1 expression. Additionally, we divided the samples into high and low LncRNA-ZMIZ1-AS1 expression groups to analyze m6A-modifying factor expression levels. Using Cibersort, we analyzed the correlation between high and low LncRNA-ZMIZ1-AS1 expression and the proportions of various immune cells in HCC. Statistical analyses of overall survival (OS) and progression-free survival (PFS) were performed using the surv_cutpoint function in the survminer package (v0.4.9), with visualization performed using ggplot2. Patients were stratified into high and low LncRNA-ZMIZ1-AS1 expression groups based on the TCGA dataset.

### Functional Enrichment Analysis

Gene Set Enrichment Analysis (GSEA) was performed to analyze KEGG pathway enrichment between high and low LncRNA-ZMIZ1-AS1 expression groups. Gene sets were uploaded to the Molecular Signatures Database in GSEA for enrichment analysis, with a false discovery rate (FDR) q-value threshold of <0.05. The top eight enriched pathways were reported.

### Cell Culture and Transfection

HepG2 cells were obtained from the American Type Culture Collection (ATCC) and cultured in Dulbecco’s modified Eagle’s medium (DMEM) supplemented with 10% fetal bovine serum at 37°C in a humidified incubator with 5% CO₂. All cell lines were routinely tested for mycoplasma and confirmed negative. The HBx overexpression lentiviral vector was purchased from Shanghai Genechem Co., Ltd. and infected into HepG2 cells using the HitransG virus transfection reagent kit. Additionally, the CAS9 Easy lentiviral (KO single vector) plasmid was purchased from Genechem and transfected into the HepG2 cells, which were previously verified and stably transfected with HBx, using Lipofectamine TM2000 to silence LncRNA-ZMIZ1-AS1. Transfection steps were performed according to the reagent instructions.

### RNA Extraction and Real-time PCR (RT-PCR)

Total RNA was extracted from cells using the Molpure® RNA extraction kit (YEASEN Co. Ltd.), and cDNA was synthesized using the PrimeScript™ RT reagent kit (TaKaRa) and stored at -20°C for later use. Real-time polymerase chain reaction (PCR) analysis was performed with the TB Green® Premix Ex Taq™ II (Tli RNaseH Plus) kit (TaKaRa), with each sample tested in triplicate. GAPDH was used as the internal reference gene. Primer sequences are provided in Table 2.

### Western Blotting

Total protein was separated by 10% SDS-PAGE and transferred onto PVDF membranes. After washing with PBS, membranes were blocked with 5% nonfat milk and incubated overnight at 4°C with primary antibodies at the manufacturer-recommended dilutions. Membranes were then incubated with secondary antibodies at room temperature for 2 hours. Immunoreactive bands were visualized using an enhanced chemiluminescence kit (Servicebio, Wuhan, China), and bands were analyzed using the Tanon Fluorescence Image Analysis System software v2.0. Band intensity was quantified using Gel Pro Analyzer software. Primary antibodies used included anti-HBx (1:1000, Abcam, Cambridge, UK), anti-PTBP1 (1:5000, Proteintech), anti-ZMIZ1 (1:1000, Zenbio), and anti-GAPDH (1:1000, Servicebio, Wuhan, China). GAPDH was used as an internal control.

### Microarray Analysis

Human m6A epitranscriptomic and mRNA microarray analysis was conducted by Arraystar (Rockville, MD, USA). Briefly, total RNA was immunoprecipitated with an anti-N6-methyladenosine (m6A) antibody, with the immunoprecipitated pellet referred to as “IP” and the recovered supernatant as “Sup.” The “IP” and “Sup” RNA were labeled with Cy5 and Cy3, respectively, hybridized to the Arraystar Human m6A Epitranscriptomic Microarray (8×60K), and scanned using an Agilent G2505C Scanner.

### MeRIP-qPCR

For MeRIP-qPCR, 1–3 μg of total RNA and an m6A control mix were combined with 2 μg of anti-m6A rabbit polyclonal antibody in 300 μL of 1×IP buffer (50 mM Tris-HCl pH 7.4, 150 mM NaCl, 0.1% NP40, and 40 U/μL RNase inhibitor). Each sample was incubated with 20 μL of Dynabeads™ M-280 sheep anti-rabbit IgG suspension, blocked with 0.5% BSA, and washed three times with 1×IP buffer before resuspension in the RNA-antibody mixture. Samples were incubated at 4°C with end-over-end rotation for 2 hours. Enriched RNA was eluted at 50°C for 1 hour and analyzed by qRT-PCR. Primer sequences are listed in Table 2.

### Transwell Migration and Invasion Assays

To assess cell migration and invasion capabilities, HepG2 cells were serum-starved for 24 hours.

An 8-μm Transwell membrane was coated with 1:8 diluted Matrigel for invasion assays (not required for migration assays). A diluted single-cell suspension (1×10^5^ cells in 200 μL) was added to the upper chamber, with 600 μL of 15% FBS medium added to the lower chamber. After 24 hours of incubation at 37°C, cells were fixed with 3.7% formaldehyde, washed with PBS, and stained with crystal violet. Cells that did not migrate/invade were removed from the upper chamber, and the number of migrated or invaded cells was counted in three independent experiments.

### Cell Viability Assay

Cell proliferation was assessed using the Cell Counting Kit-8 (CCK-8; Biosharp) following the manufacturer’s instructions. Optical density (OD) values were measured at 24, 48, and 72 hours to generate cell growth curves.

### Flow Cytometry Analysis of Apoptosis and Cell Cycle

Apoptosis was measured using the Annexin V-APC apoptosis detection kit (KeyGEN BioTECH) following the manufacturer’s protocol. After staining with 7-AAD and Annexin V/APC, apoptotic cells were analyzed using a Cytoflex flow cytometer (Beckman Coulter, Brea, CA, USA) and CytExpert software. For cell cycle analysis, cells were fixed overnight in 70% ethanol and analyzed using the KGA512 Cell Cycle Detection Kit (KeyGEN BioTECH) on a Cytoflex flow cytometer.

### Statistical Analysis

All data are expressed as mean ± standard error. Group comparisons were performed using unpaired or paired two-tailed t-tests, with the relationship between LncRNA-ZMIZ1-AS1 expression and clinicopathological features assessed via Chi-square tests. Kaplan-Meier curves were used to describe survival functions, with log-rank tests for differences between groups. Data analysis was conducted using SPSS and GraphPad software, with statistical significance set as follows: *P < 0.05, **P < 0.01, ***P < 0.001, and ****P < 0.0001.

## Results

### Association Between LncRNA-ZMIZ1-AS1 and Clinical Prognosis in HCC Patients

Using a lentiviral vector, we constructed a stable HBx-transfected HepG2 HCC cell line (HepG2-HBx) and confirmed HBx expression via Western blotting (Figure 1). Additionally, m6A epitranscriptomic microarray analysis was conducted to examine changes in non-coding RNA expression in the HBx-transfected HepG2 cell line. Bioinformatics analysis, integrating GEO and TCGA database results, indicated that LncRNA-ZMIZ1-AS1 is highly expressed in HCC cell lines and significantly associated with clinical features (Table 1).

**Figure 1:**
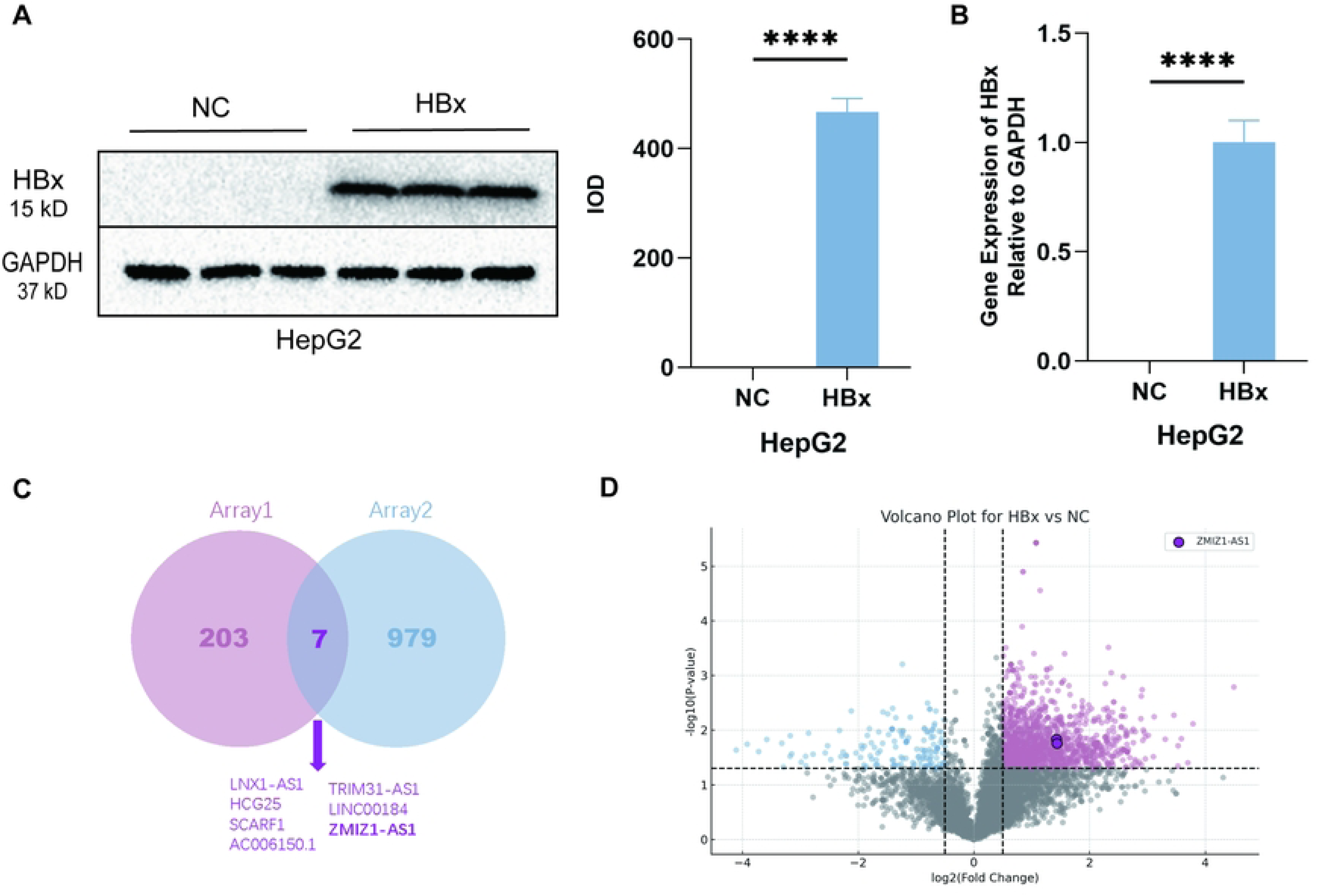
Construction of HBx-stable transfected cell lines and screening of LncRNA-ZMIZ1-AS1.

**Table 1:** Clinical characteristics of high and low LncRNA-ZMIZ1-AS1 expression groups in the TCGA dataset. No significant association was found between LncRNA-ZMIZ1-AS1 expression and age (P = 0.972), but significant relationships were observed for gender, clinical stage, and tumor stage (P = 0.005, 0.047, and 0.043, respectively).

| Clinical Information | Group |  | X2 | P Value |
| --- | --- | --- | --- | --- |
| ZMIZ1-AS1 | Low | High |  |  |
| Age |  |  | 0.001 | 0.972 |
| <65 | 112(32.7<br>%) | 111(32.5<br>%) |  |  |
| >=65 | 60(17.5%) | 59(17.3%) |  |  |
| Gender |  |  | 7.993 | 0.005 |
| Female | 67(19.6%) | 42(12.3%) |  |  |
| Male | 105(30.7<br>%) | 128(37.4<br>%) |  |  |
| Grade |  |  | 1.633 | 0.652 |
| G1 | 19(5.6%) | 26(7.6%) |  |  |
| G2 | 87(25.4%) | 81(23.7%) |  |  |
| G3 | 59(17.3%) | 58(17.0%) |  |  |
| G4 | 7(2.0%) | 5(1.5%) |  |  |
| Stage |  |  | 7.854 | 0.047 |
| S1 | 76(22.2%) | 93(27.2%) |  |  |
| S2 | 40(11.7%) | 43(12.6%) |  |  |
| S3 | 52(15.2%) | 33(9.6%) |  |  |
| S4 | 4(1.2%) | 1(0.3%) |  |  |
| T |  |  | 8.153 | 0.043 |
| T1 | 77(22.5%) | 94(27.5%) |  |  |
| T2 | 41(12.0%) | 44(12.9%) |  |  |
| T3 | 49(14.3%) | 27(7.9%) |  |  |
| T4 | 5(1.5%) | 5(1.5%) |  |  |
| N |  |  | - | 0.623 |
| N0 | 134(50.6<br>%) | 127(47.9<br>%) |  |  |
| N1 | 3(1.1%) | 1(0.4%) |  |  |
| M |  |  | - | 1 |
| M0 | 130(51.0<br>%) | 121(47.5<br>%) |  |  |
| M1 | 2(0.8%) | 2(0.8%) |  |  |

**Table 2:**
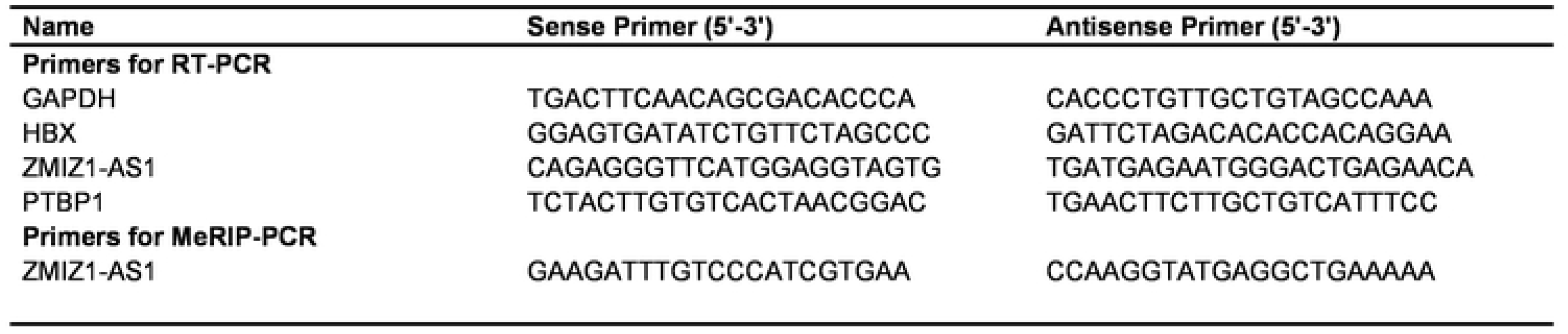
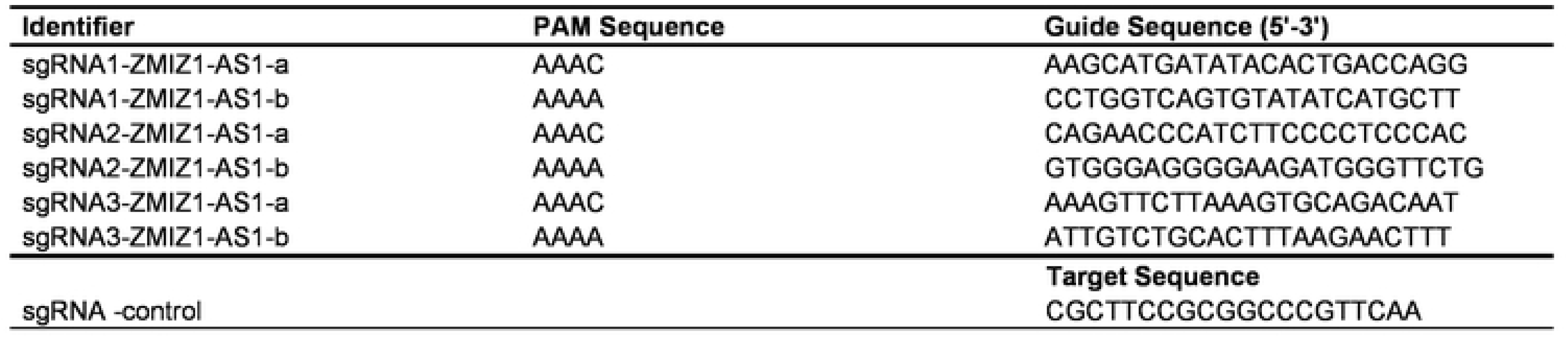
Sequences of RNA and DNA Oligonucleotides.

According to Table 1, no significant association was observed between ZMIZ1-AS1 expression and age (P = 0.972), with similar proportions in both <65 and ≥65 age groups. Gender, however, showed a significant association with ZMIZ1-AS1 expression (P = 0.005); the high-expression group included fewer females (19.6%) and more males (30.7%), indicating a higher proportion of males in the high-expression group. Tumor grade was not significantly associated with ZMIZ1-AS1 expression (P = 0.652). In contrast, clinical stage was significantly associated with ZMIZ1-AS1 expression (P = 0.047), suggesting some variation across stages. Tumor staging (T) also exhibited a significant relationship with ZMIZ1-AS1 expression (P = 0.043), indicating differences in expression proportions among groups, warranting further investigation. No significant differences were observed for lymph node and distant metastasis, with P-values of 0.623 and 1, respectively. It should be noted that the limited sample size and statistical methods may impact the results, and multiple comparisons should be taken into consideration. Larger sample sizes and additional clinical features would help improve the reliability of these conclusions.

Furthermore, m6A epitranscriptomic microarray results indicated a significant increase in m6A modification levels of LncRNA-ZMIZ1-AS1 in HBx-overexpressing cell lines, with higher expression observed compared to non-transfected HCC cells (Figure 2). Further analysis revealed that LncRNA-ZMIZ1-AS1 expression correlates with levels of m6A “reader” proteins (including Readers, Writers, and Erasers). We also observed a relationship between LncRNA-ZMIZ1-AS1 expression and immune cell infiltration (Figure 3). Based on previous experimental results, LncRNA-ZMIZ1-AS1 was selected for further analysis^[15]^.

**Figure 2:**
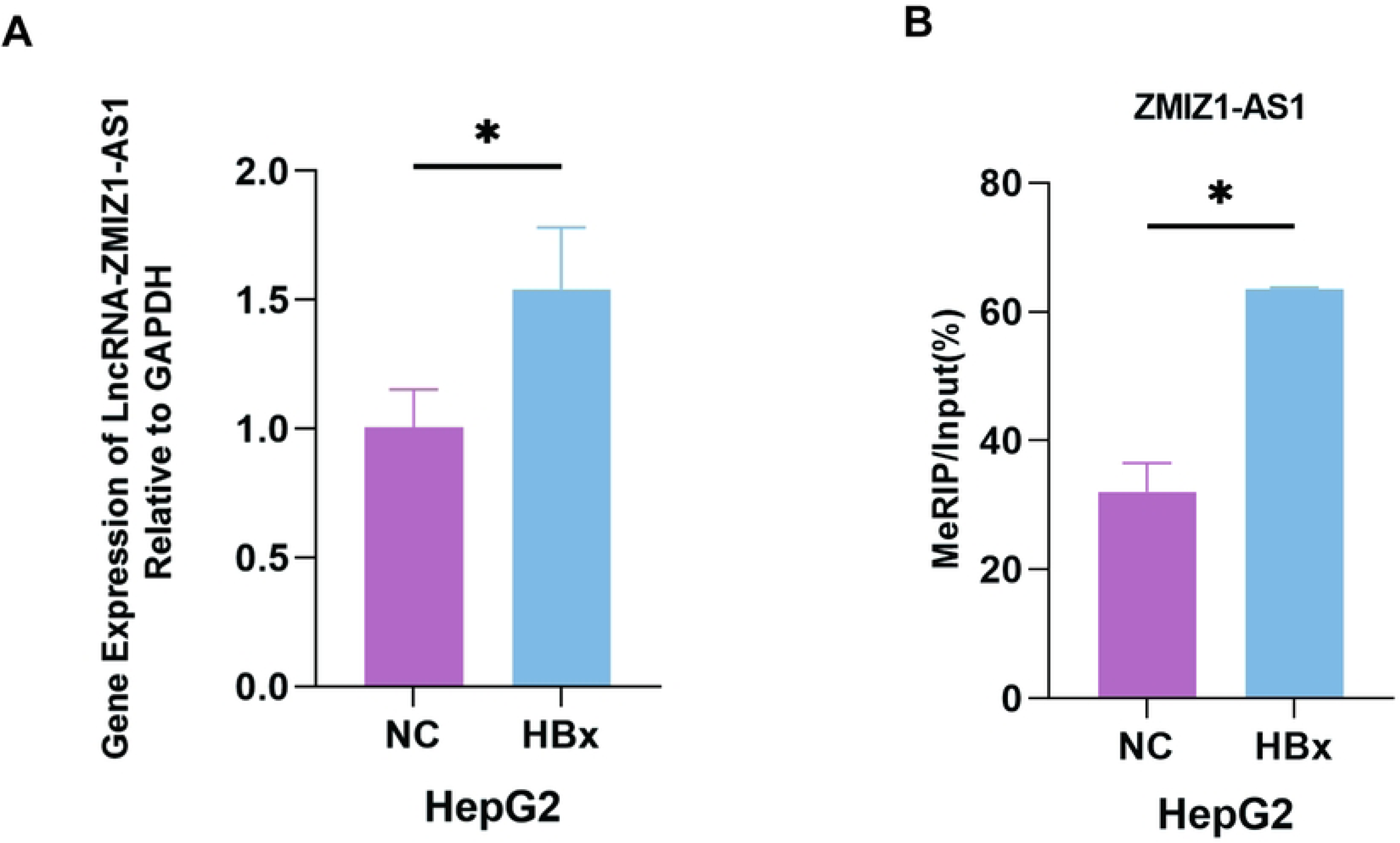
Expression and m6A modification levels of LncRNA-ZMIZ1-AS1 in HBx-stable transfected HCC cells. **A**: Data shows high expression of LncRNA-ZMIZ1-AS1 in HepG2-HBx cells. **B**: m6A modification levels of LncRNA-ZMIZ1-AS1 in HepG2-HBx cells are significantly increased. Statistical significance is indicated by *P < 0.05, **P < 0.01, ***P < 0.001, ****P < 0.0001.

**Figure 3:**
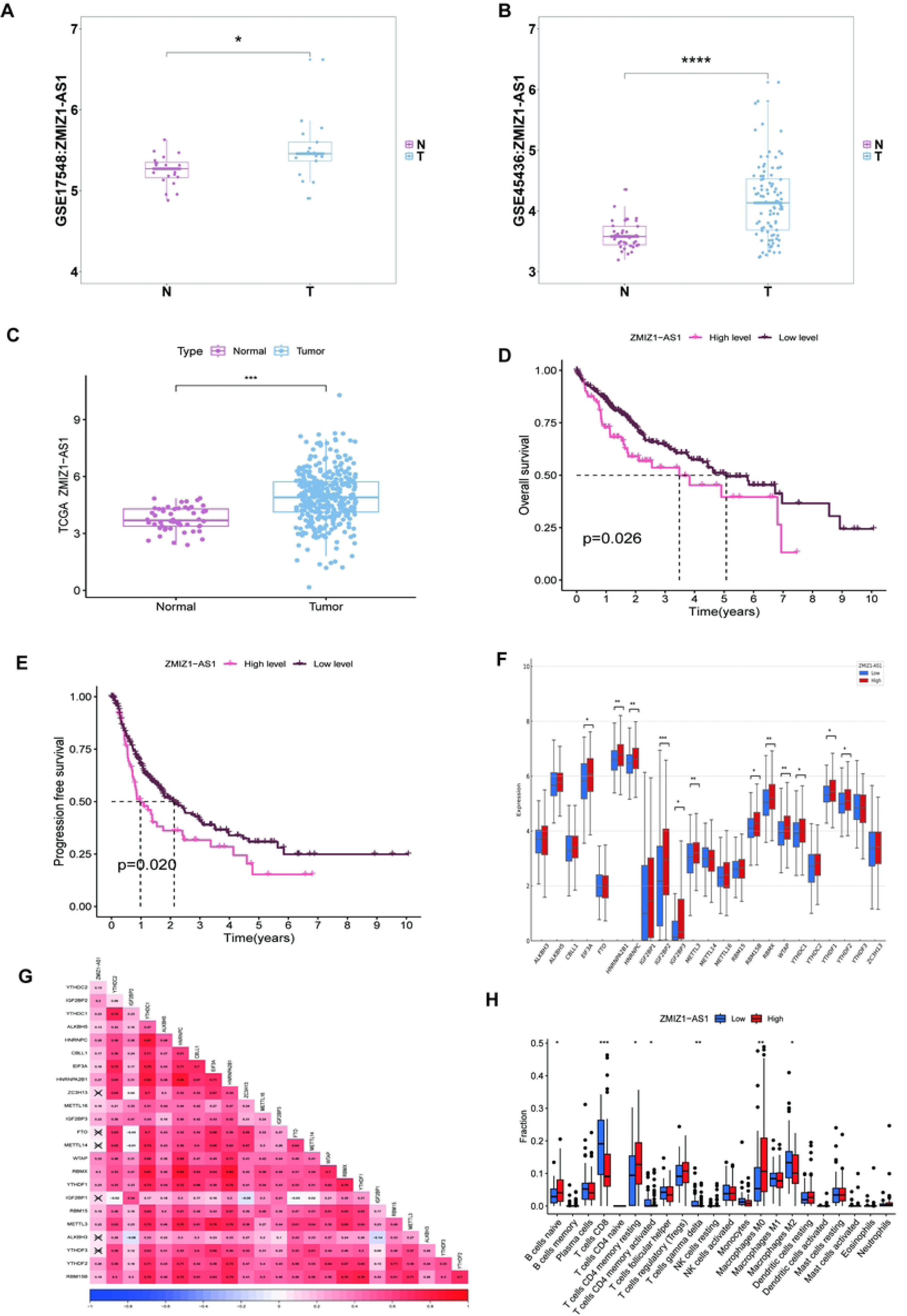
Correlation analysis of LncRNA-ZMIZ1-AS1 gene expression with clinical pathological characteristics based on GEO and TCGA databases. **A**: Expression of LncRNA-ZMIZ1-AS1 in HCC patients from the GSE17548 database is significantly higher than in non-tumor patients. **B**: GSE45436 database results show a similar trend. **C**: TCGA database further confirms the high expression of LncRNA-ZMIZ1-AS1 in HCC patients. **D - E**: Kaplan-Meier curves indicating that patients with high LncRNA-ZMIZ1-AS1 expression have shorter overall survival (OS) and progression-free survival (PFS) compared to patients with low expression. **F - G**: Differential and co-expression analysis of m6A-related genes in high and low LncRNA-ZMIZ1-AS1 expression groups. **H**: Immune infiltration plot showing the effect of LncRNA-ZMIZ1-AS1 expression levels on the infiltration of various immune cell types.

### Knockdown of LncRNA-ZMIZ1-AS1 Suppresses HCC Cell Viability, Proliferation, Migration, Invasion, and Influences Cell Cycle and Apoptosis

We designed and synthesized three sgRNAs and a negative control for ZMIZ1-AS1 and constructed sgRNA vectors to transiently knockdown ZMIZ1-AS1 expression levels (Figure 4A). The interference efficiency of lncRNA ZMIZ1-AS1 in HepG2-HBx cells was measured using qRT-PCR, and the sgRNA with the best interference effect was selected for biological behavior experiments. Using CCK-8, Transwell assays, and flow cytometry, we examined the effects of ZMIZ1-AS1 knockdown on cell proliferation, migration, invasion, and apoptosis.

**Figure 4:**
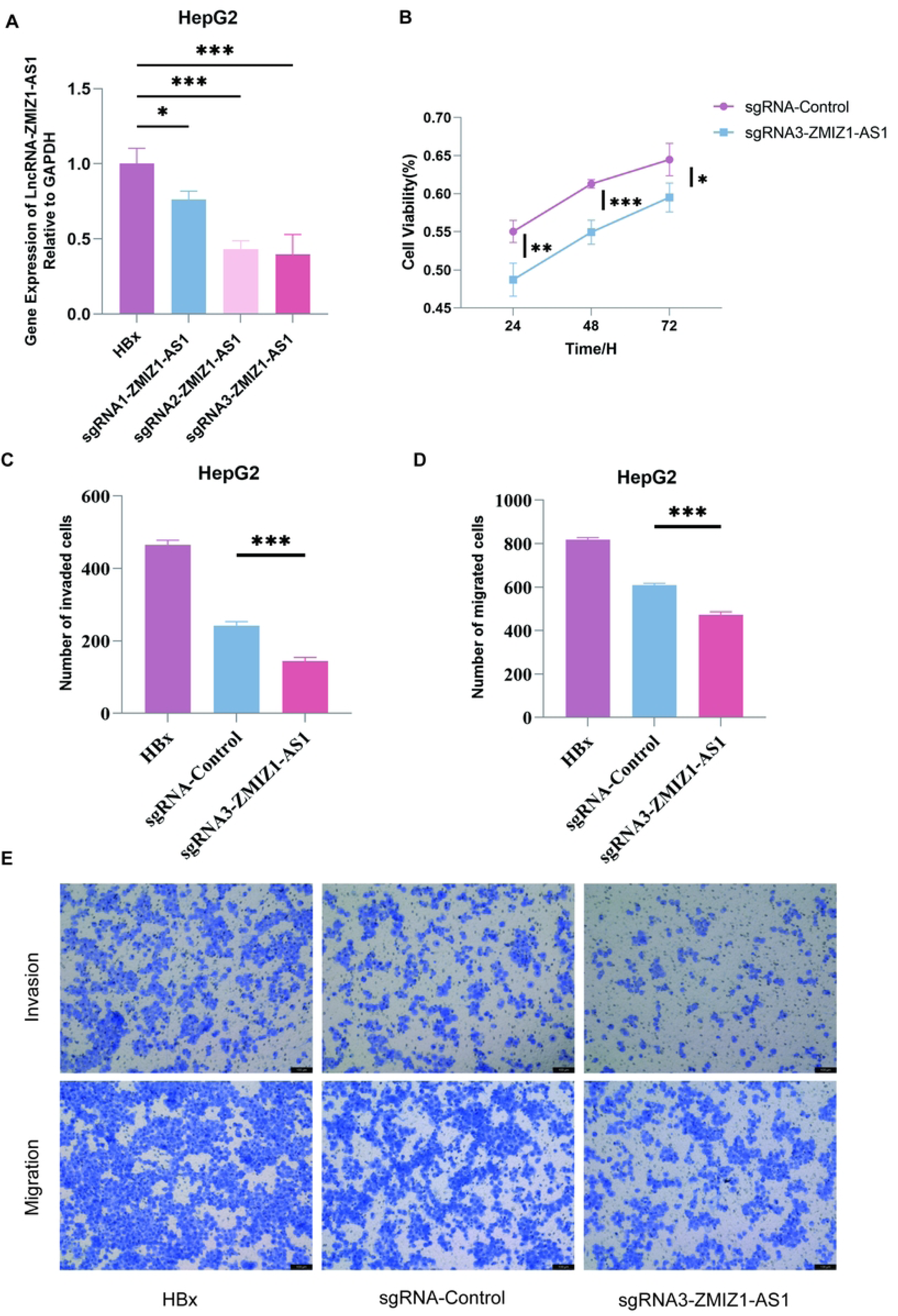
Impact of LncRNA-ZMIZ1-AS1 knockdown on cell viability, invasion, and migration. **A**: Three sgRNAs were synthesized, and qRT-PCR was used to assess interference efficiency of LncRNA-ZMIZ1-AS1 in HepG2-HBx cells, identifying the most effective sgRNA. **B**: CCK-8 assay results showing a significant decrease in cell viability at 24, 48, and 72 hours post-knockdown. **C - E**: Transwell assay results indicating that low ZMIZ1-AS1 expression significantly reduces migration and invasion capacity in HepG2-HBx cells. Statistical significance is denoted by *P < 0.05, **P < 0.01, ***P < 0.001, ****P < 0.0001.

**Figure 5:**
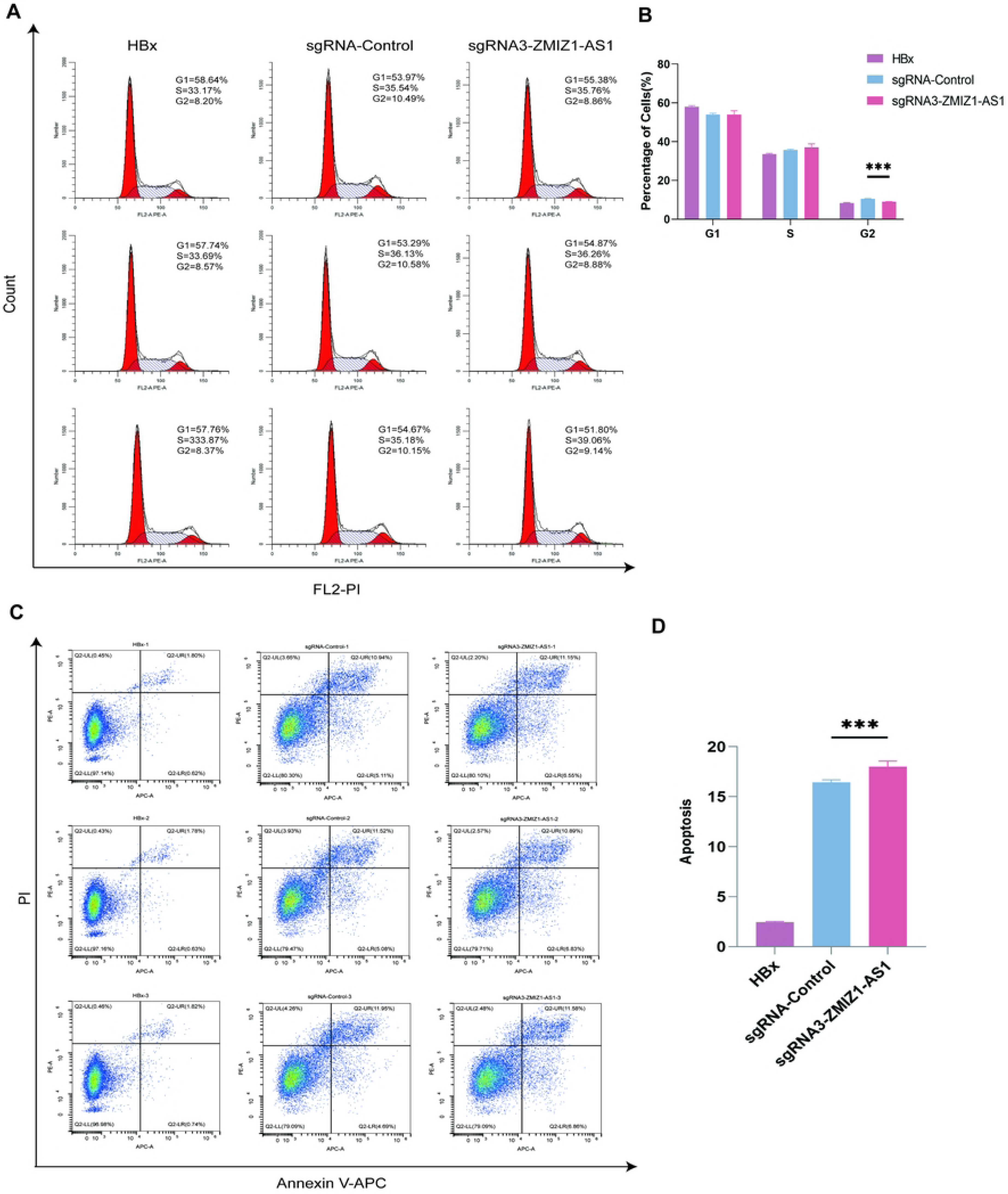
Effects of LncRNA-ZMIZ1-AS1 knockdown on cell cycle and apoptosis. **A - B**: Cell cycle analysis showing that knockdown of LncRNA-ZMIZ1-AS1 significantly reduces the proportion of cells in the G2 phase (n=3, t-test). **C - D**: Apoptosis assay results showing that low ZMIZ1-AS1 expression significantly increases apoptosis in HepG2-HBx cells (n=3, t-test). Statistical significance is marked as *P < 0.05, **P < 0.01, ***P < 0.001, ****P < 0.0001. **A - B**: Western blot analysis showing HBx expression levels in HepG2 cell lines, with significant expression observed in the stable transfected group. **C**: Venn diagram showing the intersection of two microarrays, including candidate genes such as LncRNA-ZMIZ1-AS1, HCG25, TRIM31-AS1, LINC00184, SCARF1, and AC006150.1. **D**: Volcano plot showing differential expression in the m6A epitranscriptomic microarray. Significant genes are marked as *P < 0.05, **P < 0.01, ***P < 0.001, ****P < 0.0001.

Using the sgRNA3 sequence with optimal interference efficiency, we transiently transfected HepG2-HBx cells to knock down lncRNA ZMIZ1-AS1, creating three groups: blank control, sgRNA3-ZMIZ1-AS1, and sgRNA-control. CCK-8 assays were performed in 96-well plates to assess the optical density (OD) at 24h, 48h, and 72h, confirming the effect of lncRNA ZMIZ1-AS1 on cell proliferation. Results showed a statistically significant reduction in cell activity at 24h, 48h, and 72h in the knockdown group (Figure 4B).

Similarly, we performed Transwell migration and invasion assays by counting the number of HCC cells invading the Matrigel 24 hours post-transfection. The number of cells invading the matrix gel was significantly higher in the sgRNA-control group than in the sgRNA3-ZMIZ1-AS1 group, with a statistically significant difference between the two groups (Figures 4C–E). These results indicate that ZMIZ1-AS1 expression affects not only the migration capacity of HCC cells but also their invasion ability.

### Positive Regulation of ZMIZ1 Expression by LncRNA-ZMIZ1-AS1 and Potential Involvement with the NOTCH Signaling Pathway

Previous studies have shown that lncRNA ZMIZ1-AS1 interacts with both ZMIZ1 and PTBP1. To investigate this mechanism, we constructed lncRNA ZMIZ1-AS1 knockdown and control groups using transient transfection and assessed PTBP1 and ZMIZ1 expression levels via Western blotting. Results demonstrated that knockdown of lncRNA-ZMIZ1-AS1 significantly reduced ZMIZ1 expression levels but had no significant effect on PTBP1 expression (Figure 6). This suggests that LncRNA-ZMIZ1-AS1 may stabilize ZMIZ1 expression through interaction with PTBP1, thereby promoting the malignant phenotype in tumor cells.

**Figure 6:**
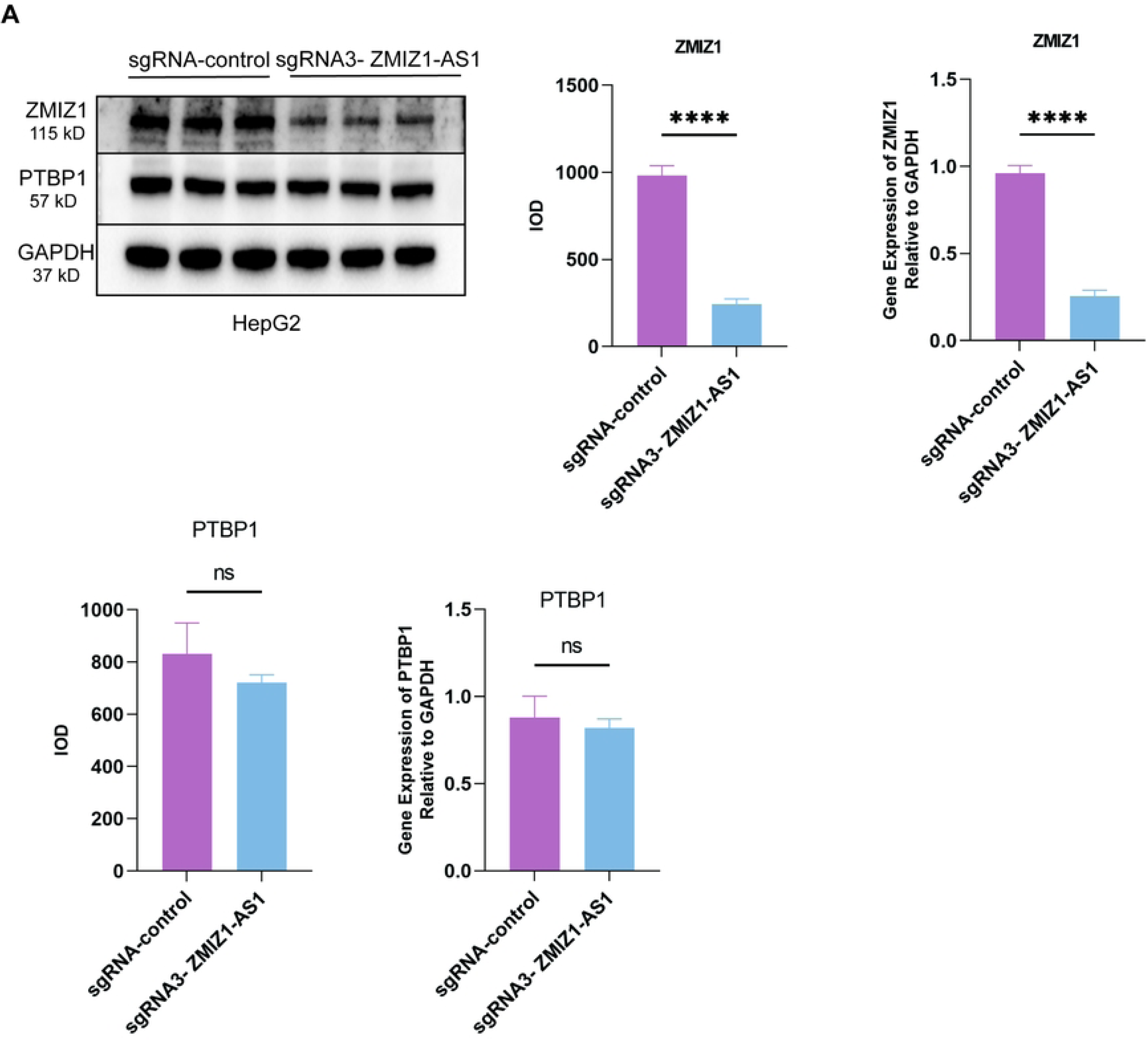
Regulatory effects of LncRNA-ZMIZ1-AS1 on PTBP1 and ZMIZ1 expression levels. **A**: Western blot analysis showing no significant difference in PTBP1 expression between LncRNA-ZMIZ1-AS1 knockdown and control groups (n=3, t-test). **B**: ZMIZ1 expression is significantly reduced in the LncRNA-ZMIZ1-AS1 knockdown group, suggesting that LncRNA-ZMIZ1-AS1 may regulate ZMIZ1 stability through interaction with PTBP1 (n=3, t-test). Statistical significance is marked as *P < 0.05, **P < 0.01, ***P < 0.001, ****P < 0.0001.

Through bioinformatics and GSEA analyses, we further explored differences between high and low LncRNA-ZMIZ1-AS1 expression groups. KEGG enrichment analysis indicated that LncRNA-ZMIZ1-AS1 may be associated with several signaling pathways, including NOTCH, ERBB, NEUROTROPHIN, INSULIN, GNRH, VEGF, and TOLL LIKE RECEPTOR pathways (Figure 7). These pathways play significant roles in tumorigenesis and progression, suggesting that LncRNA-ZMIZ1-AS1 may modulate HCC cell proliferation, migration, and apoptosis via pathways such as NOTCH.

**Figure 7:**
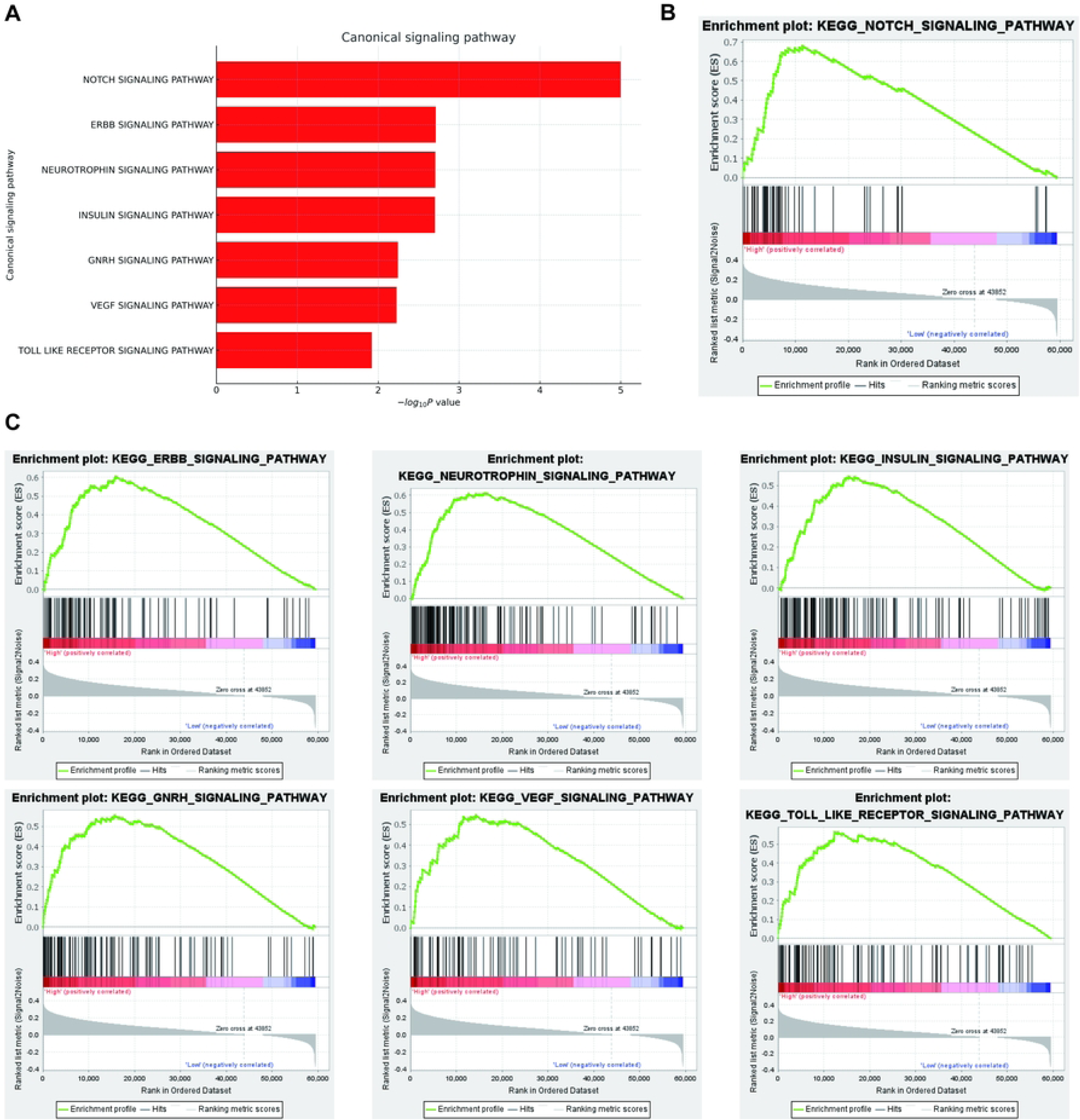
Enrichment analysis of genes co-expressed with LncRNA-ZMIZ1-AS1. **A**: GSEA enrichment analysis showing significant enrichment of co-expressed genes in the top eight signaling pathways. **B**: KEGG enrichment analysis indicating a correlation with the NOTCH signaling pathway. **C**: Other associated pathways, including ERBB, NEUROTROPHIN, INSULIN, GNRH, VEGF, and TOLL LIKE RECEPTOR pathways.

## Discussion

The fundamental mechanism of the Notch signaling pathway involves the binding of Notch receptors to ligands (such as DLL1, DLL3, DLL4, JAG1, and JAG2) on adjacent cells. This interaction triggers a series of proteolytic cleavages by enzymes, including ADAM and γ-secretase, which release the Notch intracellular domain (NICD) of the receptor. NICD then enters the nucleus, where it binds to co-activators such as RBPJ (CSL) and MAML to form the Notch transcriptional activation complex (NTC), which activates downstream genes. Notch signaling not only directly activates or represses target gene transcription but also finely tunes gene expression through chromatin remodeling mechanisms such as histone modifications. Consequently, this pathway impacts processes like cell differentiation and cancer progression. In tumor biology, Notch signaling functions in both tumor cells and the tumor microenvironment, where it modulates immune cells, indirectly influencing tumor growth and immune evasion. Notch signaling plays a dual role in regulating the function of immune cells within the tumor microenvironment, including tumor-associated macrophages (TAMs), myeloid-derived suppressor cells, dendritic cells, and T cells, in the context of antitumor or pro-tumor immune responses^[16, 17]^.

Both bioinformatics and functional experiments indicate that high expression of LncRNA-ZMIZ1-AS1 in HCC cells is closely associated with enhanced proliferation, migration, and invasion capabilities. Additionally, when analyzing the correlation between LncRNA-ZMIZ1-AS1 and clinical-pathological characteristics, significant differences were found in gender, clinical stage, and tumor staging, suggesting that LncRNA-ZMIZ1-AS1 may serve as a potential biomarker for HCC progression. These findings align with previous research that observed LncRNA-ZMIZ1-AS1 promoting malignant phenotypes in other types of cancers.

M6A epitranscriptomic microarray analysis revealed significant m6A modification of LncRNA-ZMIZ1-AS1 in HBx-stably transfected HepG2 cells, suggesting that m6A modification may influence the stability, transcription, or translation efficiency of LncRNA-ZMIZ1-AS1 in HCC cells. In recent years, m6A modification has emerged as a critical RNA modification in cancer^[18, 19]^. The m6A modification of LncRNA-ZMIZ1-AS1 may enable it to bind specific proteins or RNAs, thereby influencing tumor occurrence and progression. Further analysis indicated that high expression of LncRNA-ZMIZ1-AS1 was significantly associated with the levels of m6A “reader” proteins, which may regulate its function by recognizing m6A-marked LncRNA-ZMIZ1-AS1. Future research should explore how different m6A modification patterns impact LncRNA-ZMIZ1-AS1 function to uncover its role in cancer at the epigenetic level.

Knockdown of LncRNA-ZMIZ1-AS1 significantly reduced HCC cell viability, proliferation, migration, and invasion capabilities. Transwell and flow cytometry assays demonstrated that LncRNA-ZMIZ1-AS1 knockdown not only inhibits cell migration but also decreases the ability to invade the extracellular matrix, suggesting that LncRNA-ZMIZ1-AS1 may influence metastatic potential by regulating relevant signaling pathways. Additionally, cell proliferation assays indicated that LncRNA-ZMIZ1-AS1 may promote HCC cell proliferation by regulating cell cycle-related pathways or factors, warranting further investigation into its molecular mechanisms. In this study, knockdown of LncRNA-ZMIZ1-AS1 significantly increased apoptosis in HCC cells, suggesting its role in negatively regulating apoptosis. This finding aligns with previous research indicating that LncRNA-ZMIZ1-AS1 may suppress apoptosis in tumor cells by interacting with other apoptosis-related molecules or pathways, thus promoting cell survival. Bioinformatics analysis shows that LncRNA-ZMIZ1-AS1 is associated with multiple tumor-related signaling pathways, such as the NOTCH pathway, providing a foundation for further exploration of its molecular mechanisms in apoptosis regulation.

Our results demonstrate that LncRNA-ZMIZ1-AS1 positively regulates ZMIZ1 expression but does not significantly impact PTBP1 expression. This suggests that LncRNA-ZMIZ1-AS1 may stabilize ZMIZ1 expression by interacting with PTBP1, thereby promoting malignant phenotypes in tumor cells. Previous research also suggested a regulatory interaction between LncRNA-ZMIZ1-AS1 and PTBP1, reinforcing its role in the regulation of neighboring gene expression. The high expression of LncRNA-ZMIZ1-AS1 in HCC and its promotion of malignant phenotypes imply its potential as a diagnostic and therapeutic biomarker for HCC. The Notch signaling pathway is a highly conserved cell communication mechanism that plays an essential regulatory role in multicellular organisms. By regulating processes like cell proliferation, differentiation, and apoptosis, Notch signaling is involved in embryonic development, adult tissue homeostasis, and cellular behavior in various pathological states. Future research could focus on its role in the tumor microenvironment and immune evasion to explore the potential of LncRNA-ZMIZ1-AS1 in HCC immunotherapy. Additionally, m6A modification may be a critical aspect of LncRNA-ZMIZ1-AS1 regulation, providing new perspectives and directions for epigenetic research in cancer. In-depth studies of m6A modification in conjunction with other signaling pathways, such as NOTCH, may reveal more about the regulation and function of LncRNA-ZMIZ1-AS1.

It is important to note that while this study has achieved certain insights into the role of LncRNA-ZMIZ1-AS1 in regulating malignant phenotypes in HCC cells, there are limitations. First, detailed pathway mechanism studies are lacking, such as experiments investigating how LncRNA-ZMIZ1-AS1 regulates HCC cell behavior through specific signaling pathways (e.g., NOTCH or other tumor-related pathways). This limits our understanding of its molecular mechanisms. Additionally, no rescue experiments were conducted to explore the interactions between LncRNA-ZMIZ1-AS1, PTBP1, and ZMIZ1. Conducting rescue experiments to determine whether exogenous expression of PTBP1 or ZMIZ1 can restore partial malignant phenotypes in HCC cells after LncRNA-ZMIZ1-AS1 knockdown would more effectively validate the role of PTBP1 in this regulatory mechanism. Furthermore, in vivo studies using mouse models were not performed to verify the specific role of LncRNA-ZMIZ1-AS1 in HCC progression. Although in vitro cell experiments have revealed the impact of LncRNA-ZMIZ1-AS1 on HCC cell proliferation, migration, and invasion, the lack of in vivo models limits the ability to fully reflect its potential influence within the tumor microenvironment.

In summary, our study preliminarily demonstrates that LncRNA-ZMIZ1-AS1 plays an essential role in promoting malignant phenotypes in HCC cells, as its high expression facilitates cell proliferation, migration, and invasion. While limitations exist in pathway mechanism studies, rescue experiments, and in vivo validation, our findings provide initial evidence of the biological function of LncRNA-ZMIZ1-AS1 in HCC, laying the groundwork for future exploration of its regulatory mechanisms. Additionally, the experimental results support the potential role of LncRNA-ZMIZ1-AS1 in cancer epigenetic regulation, particularly in m6A modification and PTBP1 interaction, warranting further investigation.

## Authorship

The authors confirm contribution to the paper as follows: study conception and design: Gang Wu;data collection: Hao Tang;analysis and interpretation of results: Chunhu Mao, Guo Chen;draft manuscript preparation: Gang Wu.All authors reviewed the results and approved the final version of the manuscript.

## Ethics Approval and Informed Consent Statement

None

## Availability of Data and Materials

All authors can guarantee the authenticity and usability of the data and materials in the article. All the data used in this study can be found in the main paper. The datasets will be available from the corresponding author on reasonable request.

## Reference

[1] De Lope C R, Tremosini S, Forner A, et al. Management of HCC [J]. Journal of hepatology, 2012, 56 Suppl 1: S75–87.

[2] Cauchy F, Soubrane O, Belghiti J. Liver resection for HCC: patient’s selection and controversial scenarios [J]. Best practice & research Clinical gastroenterology, 2014, 28(5): 881–96.

[3] Thabut D, Kudo M. Treatment of portal hypertension in patients with HCC in the era of Baveno VII [J]. Journal of hepatology, 2023, 78(3): 658–62.

[4] Brown Z J, Tsilimigras D I, Ruff S M, et al. Management of Hepatocellular Carcinoma: A Review [J]. JAMA surgery, 2023, 158(4): 410–20.

[5] Wang J, Zhu S, Meng N, et al. ncRNA-Encoded Peptides or Proteins and Cancer [J]. Molecular therapy : the journal of the American Society of Gene Therapy, 2019, 27(10): 1718–25.

[6] Nemeth K, Bayraktar R, Ferracin M, et al. Non-coding RNAs in disease: from mechanisms to therapeutics [J]. Nature reviews Genetics, 2024, 25(3): 211–32.

[7] Slack F J, Chinnaiyan A M. The Role of Non-coding RNAs in Oncology [J]. Cell, 2019, 179(5): 1033–55.

[8] Tan Y T, Lin J F, Li T, et al. LncRNA-mediated posttranslational modifications and reprogramming of energy metabolism in cancer [J]. Cancer communications (London, England), 2021, 41(2): 109–20.

[9] Bridges M C, Daulagala A C, Kourtidis A. LNCcation: lncRNA localization and function [J]. The Journal of cell biology, 2021, 220(2).

[10] Zhou Y, Jin Q, Chang J, et al. Long non-coding RNA ZMIZ1-AS1 promotes osteosarcoma progression by stabilization of ZMIZ1 [J]. Cell biology and toxicology, 2022, 38(6): 1013–26.

[11] Li X, Wu A, Wang Y, et al. Knockdown of circZMIZ1 enhances the anti-tumor activity of CD8(+) T cells to alleviate hepatocellular carcinoma [J]. Functional & integrative genomics, 2024, 24(1): 27.

[12] Jiang H, Lv D J, Song X L, et al. Upregulated circZMIZ1 promotes the proliferation of prostate cancer cells and is a valuable marker in plasma [J]. Neoplasma, 2020, 67(1): 68–77.

[13] Wang H W, Hsieh T H, Huang S Y, et al. Forfeited hepatogenesis program and increased embryonic stem cell traits in young hepatocellular carcinoma (HCC) comparing to elderly HCC [J]. BMC genomics, 2013, 14: 736.

[14] Yildiz G, Arslan-Ergul A, Bagislar S, et al. Genome-wide transcriptional reorganization associated with senescence-to-immortality switch during human hepatocellular carcinogenesis [J]. PLoS One, 2013, 8(5): e64016.

[15] Wu G, Yu F, Xiao Z, et al. Hepatitis B virus X protein downregulates expression of the miR-16 family in malignant hepatocytes in vitro [J]. British journal of cancer, 2011, 105(1): 146–53.

[16] Li X, Yan X, Wang Y, et al. The Notch signaling pathway: a potential target for cancer immunotherapy [J]. Journal of hematology & oncology, 2023, 16(1): 45.

[17] Wang H, Zang C, Liu X S, et al. The role of Notch receptors in transcriptional regulation [J]. Journal of cellular physiology, 2015, 230(5): 982–8.

[18] Sun T, Wu R, Ming L. The role of m6A RNA methylation in cancer [J]. Biomedicine & pharmacotherapy = Biomedecine & pharmacotherapie, 2019, 112: 108613.

[19] An Y, Duan H. The role of m6A RNA methylation in cancer metabolism [J]. Molecular cancer, 2022, 21(1): 14.

